# Development of a new genotype–phenotype linked antibody screening system

**DOI:** 10.1101/2024.03.06.581777

**Authors:** Takashi Watanabe, Hikaru Hata, Yoshiki Mochizuki, Fumie Yokoyama, Tomoko Hasegawa, Naveen Kumar, Tomohiro Kurosaki, Osamu Ohara, Hidehiro Fukuyama

## Abstract

Antibodies are powerful tools for the therapy and diagnosis of various diseases. In addition to conventional hybridoma-based screening, recombinant antibody-based screening has become a common choice; however, its application is hampered by two factors: 1) screening starts after Ig gene cloning and recombinant antibody production only, and 2) the antibody is composed of paired chains, heavy and light, commonly expressed by two independent expression vectors. Here, we introduce a method for the rapid screening of recombinant monoclonal antibodies by establishing a Golden Gate-based dual-expression vector and *in-vivo* expression of membrane-bound antibodies. Using this system, we demonstrate the rapid isolation of influenza cross-reactive antibodies with high affinity from immunized mice within 7 days. This system is particularly useful for isolating therapeutic or diagnostic antibodies, e.g., during foreseen pandemics.

**Impact Statement:** A Golden Gate-based dual-expression vector enables rapid screening, which facilitates efficient isolation of high-affinity cross-reactive antibodies for therapeutic or diagnostic use and provides a crucial advance for pandemic preparedness.

## Introduction

Technical improvements in the rapid isolation of monoclonal antibodies (mAbs) are critical for the diagnostic and therapeutic development of such antibodies. This need became apparent to the general public during the COVID-19 pandemic. Antibodies isolated from convalescent COVID-19 patients are valuable resources for the development of mAb therapeutics and rapid diagnostic tools. Human antibodies can be administered to other patients without the need to humanize in an emergency and with a slight modification of Fc receptor binding (1) to increase the half-life or reduce the risk of antibody-dependent enhancement. Although production costs remain high, the efficiency of generating recombinant antibodies has dramatically improved with the use of mammalian cell lines, and the application of therapeutic antibodies is likely to expand in the near future (2).

The gold standard method for isolating mAbs is the hybridoma technology developed by Kohler and Milstein in 1975 (3), in which antibody-producing B cells are fused with immortal B cells, called myelomas, to produce long-lasting antibody-producing B cells. In addition to hybridoma technology, new methods have been developed to increase mAb screening efficiency. These include direct immortalization of B cells by gene reprogramming using the Epstein–Barr virus (4) or retrovirus-mediated gene transfer (5), cloning of variable region-encoding genes by single-cell PCR (6,7), single-cell culture screening (8), and *in-vitro* screening of recombinant antibody libraries (9–13). Some of these methods have successfully yielded high-affinity antibodies in various formats, including single variable domain on a heavy chain or single-chain fragment variable antibodies. Although hybridoma technology is highly reliable for the isolation of valuable antibodies from animal models, such as mice, rats, and hamsters, it is time-consuming and requires technical skill. The development of a new fusion partner cell line, SPYMEG, has enabled the production of human hybridomas and opened a new direction for the isolation of human mAbs (14). Although many of these new technologies have increased the throughput for mAb isolation, the procedure still requires significant resources and time.

To dramatically enhance the efficiency of mAb isolation, we used next-generation sequencing (NGS) technology, which has revolutionized the sequencing of immunoglobulin (Ig) variable-region genes. For instance, tens of thousands of Ig genes specific to certain antigens can be identified by combining droplet-based single-cell isolation with DNA barcode antigen technology, followed by NGS (15). Although Ig genes can be sequenced at high throughput, there is no method for screening antibodies in a high-throughput format that is compatible with NGS technology.

In this study, we developed a new functional screening method that is compatible with NGS to rapidly identify antigen-specific clones. We first generated an Ig dual-expression vector using Golden Gate Cloning (16), which enabled the linkage of heavy-chain variable and light-chain variable DNA fragments obtained from a single-sorted B cell, followed by the expression of membrane-bound Ig. This single-step procedure enabled the enrichment of antigen-specific, high-affinity Igs by flow cytometry, which is significantly faster than conventional cloning-based methods that require sequential steps. To demonstrate the efficiency of our new method, we screened for potent broadly reactive antibodies (17–19) against the influenza virus in an experimental mouse model using our technology. Broad reactivity antibodies allow for the development of influenza vaccines that are effective across seasonal variations in influenza strains, which can only be accomplished by screening a large number of candidate antibodies. We first raised cross-reactive B cells against various hemagglutinin (HA) antigens from the influenza virus by sequential immunization with heterotypic HA antigens from group 1 influenza. Using this technology, we obtained several mAbs that bind to other group 1 HA antigens, and even to group 2 HA antigens, of the influenza virus. Our technology can also be applied to human antibody screening and represents a new line of mAb screening that accelerates the isolation of therapeutic and diagnostic mAbs.

## Results

### Rapid membrane-bound dual Ig expression screening system

Cloning-based Ig screening involves several steps: cloning two Ig gene fragments that encode heavy and light chains independently from a single cell, co-expressing these Ig chains, and purifying individual recombinant antibodies. These steps are labor intensive. We established a new antibody screening system for the isolation of valuable mAbs (details in the methods). Eventually, the antigen-binding Ig transformants were collected by sorting in a bulk fashion, and the unique CDR3 region and clones of interest were identified using the Ig-seq database (Fig. 1A).

**Figure 1.**
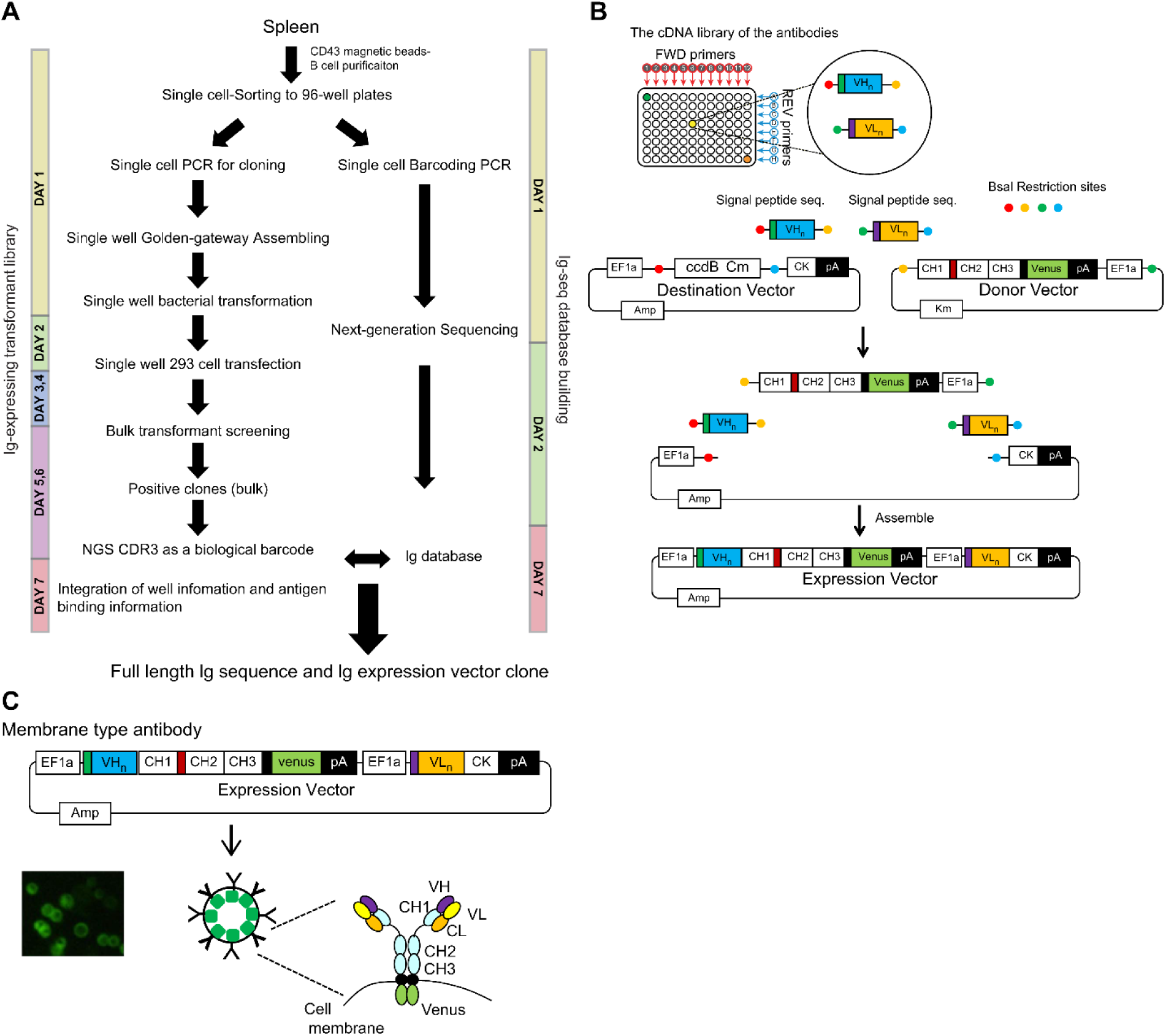
System to link the antigen-binding feature with the Ig repertoire genetic information. **A.** Single cell-sorted B cells were subjected to two processes: Ig-seq database building and Ig-expressing transformant library preparation. Antigen-binding Ig transformants were collected by sorting in bulk and the sequences of the unique CDR3 regions and clones of interest were identified by referring to the Ig-seq database. The duration of each steps are indicated. **B.** To express both Ig heavy and kappa/lambda chains in a single-expression vector, we generated a dual-expression vector. Four gene fragments were assembled by the Golden Gate method using the BsaI restriction enzyme. They included: 1) the destination vector containing the Ig kappa constant gene and the ccdB Chloramphenicol cassette for negative selection, 2) fragments containing the Ig heavy constant gene fused to *Venus* derived from the donor vector, 3) the Ig heavy variable gene fragment, and 4) the Ig light variable gene fragment. **C.** Individually purified plasmids were transiently transfected to the floating human FreeStyle 293 cell line. Igs were expressed on the cell surface in 2 days, and expression levels were confirmed and normalized by *Venus* expression, since the Ig heavy chain was fused to *Venus* at the cytoplasmic domain tail.

We generated a dual-expression vector to express both Ig heavy and kappa/lambda in a single-expression vector (Fig. 1B). These plasmids were individually transfected transiently to the floating human FreeStyle 293 cell line. Igs were expressed on the cell surface within 2 days, and their expression could be confirmed and normalized by *Venus* (20) expression since the Ig heavy chain is fused with *Venus* at the cytoplasmic domain tail (Fig. 1C). The advantage of this system is the rapid enrichment of clones of interest through bulk screening. In the current single-cell-based cloning/screening, plasmid DNA extracted from the collected antigen-binding bulk transformants is sequenced for the heavy chain CDR3 region. The CDR3 region is unique and can be used for the identification of a clone (21,22). This membrane-bound Ig expression system links antigen-binding features and genetic information of the Ig repertoire.

### Building the Ig-seq database and analysis

As a model experiment, we obtained broadly reactive antibodies against influenza viruses using multiple HA probes. We prepared two HA proteins as probes: A/Puerto Rico/8/1934 (H1N1), designated as PR8, and A/Okuda/1957 (H2N2), designated as H2. A total of 374 IgG1^+^ B cells of either PR8^+^ (204 cells), H2^+^ (99 cells), or PR8^+^H2^+^ (71 cells) were collected in a single-cell fashion. We obtained sequences of 284 independent clones. The success rate of cloning the paired Ig fragments was 75.9%. An overview of heavy chain V-D-J and light chain V-J usage and repertoire clonality is shown in Supplementary Fig. 1A and B. Mutation rates and CDR3 lengths of the heavy chains of the three cell populations (PR8^+^, H2^+^, and PR8^+^H2^+^) were comparable (Supplementary Fig. 1C and D). These results indicate that broadly reactive antibodies do not require unique genetic traces to obtain breadth in our experimental setup.

### Isolation of H2-and H1-reactive B cells

On day 2 after transfection of individual 284 clones, the transformants were mixed and stained with HA probes (Fig. 2A). As shown in Fig. 2B, we selected three prominent populations, H1^+^H2^+^ (cross), H2^+^ (H2), and H1^+^ (H1), and collected them in bulk. In contrast to bacterial transfection, mammalian cell lines, such as FreeStyle 293 cells, can contain multiple plasmids in a cell and ectopically express multiple proteins. Therefore, we decided to mix the transformants instead of plasmids. We sorted a total of 2,981 cells, classified into three strong binder populations, and performed sequencing (Fig. 2B). These bulk Ig-seq data are referred to as Ig-seq data, as described above. A total of 190 clones from the three populations were verified for their sequences and HA binding by flow cytometry analysis, as shown in Fig. 2C; 110 “H1^+^,” 67 “H2^+^,” and 13 “cross” clones were successfully isolated. In parallel, all the transformants were examined individually for comparison with the bulk method. One hundred thirty-eight transformants were verified for their binding to HAs by flow cytometry. Of these, 81 “H1^+^,” 48 “H2^+^,” and 9 “cross” clones were successfully isolated (Fig. 2D). Eight of the 13 “cross” clones obtained in the bulk examination overlapped with “cross” clones obtained in the individual examination (Fig. 2D and E); the binding of one cross clone (D11p4) from the individual examination was so weak that it was not included in the gates sorted in bulk (Fig. 2B). Comparison of the bulk and the individual examinations demonstrated that the rapid bulk method is sufficient for selecting clones of interest.

**Figure 2.**
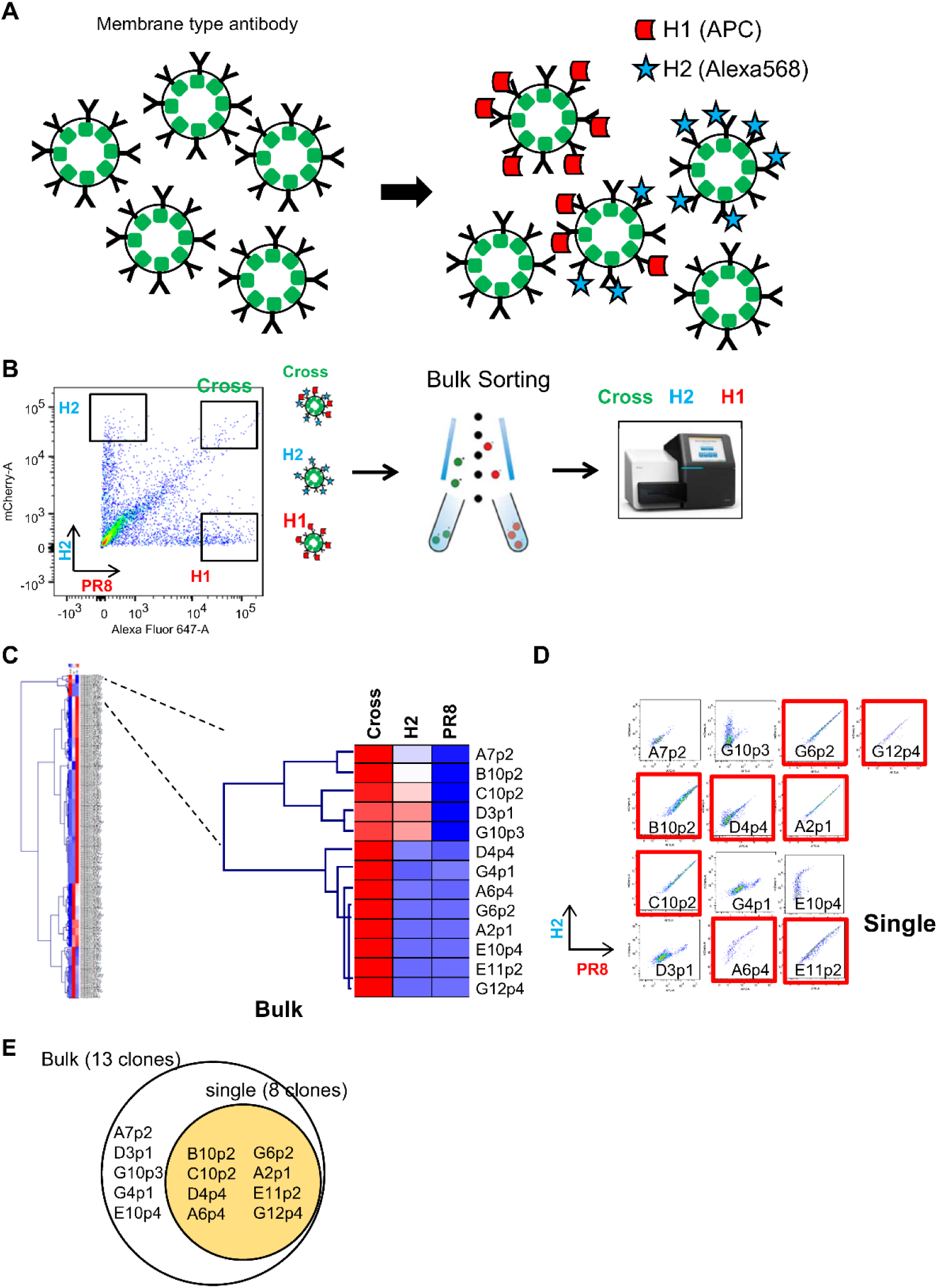
Isolation of H2 and H1-reactive B cells. **A.** Mixture of Ig-expressing transformant libraries stained with HA probes. **B.** Three strong HA-binding populations, H1^+^H2^+^ (cross), H2^+^ (H2), and H1^+^ (H1), were sorted and collected in a bulk fashion. Then, these three collected populations were and sequenced. **C.** The bulk Ig-seq data were referred to as the single-cell Ig-seq data. Heatmap indicates the appearance rate of individual clone in each box shown in B (H1, H2, or Cross) among sorted 2,981 cells of mixture of 190 clone transformants. H1^+^H2^+^ (cross) clones are shown on the right columns. High to low appearance rate reflects red to blue color. **D.** The individual flow cytometry profile of 13 “Cross” transformants obtained by the bulk examination are shown and red labeled ones are also found by the individual examination(red). **E.** A summary of the results in comparison of the bulk and individual examination methods.

### Characterization of “cross” clones for broad reactivity against the influenza virus

We decided to further characterize eight common clones between the individual and bulk examinations and one short-ranged clone, D11p4. By generating a secretory form of mAbs for these nine “cross” clones from the individual examination, we characterized their affinity and broadness using six different HA antigens from strains A/Okuda/1957 (H2N2) named for H2, A/Puerto Rico/8/1934 (H1N1) for PR8, A/California/2009 (X-179A) [H1N1] Pdm09 for Cal, A/Texas/50/2012 (X-223) [H3N2] for H3, A/Egypt/N03072/2010(H5N1) for H5, and A/Brisbane/59/2007 [H1N1] (19) for Stem. We determined the affinities (K_d_) of the antibodies for the six HA antigens using surface plasmon resonance analysis (Table 1 and Supplementary Fig. 2), which ranged from 500 to 100 nM. With the highest affinity among them, the C10p2 antibody bound to Cal at K_d_ ≃ 5.66 × 10^­10^ (M). Surprisingly, the antibodies showed a broader spectrum of HA binding among the influenza strains. Seven of the nine mAbs bound to HA from the highly pathogenic avian influenza strain H5N1.

**Table 1.**
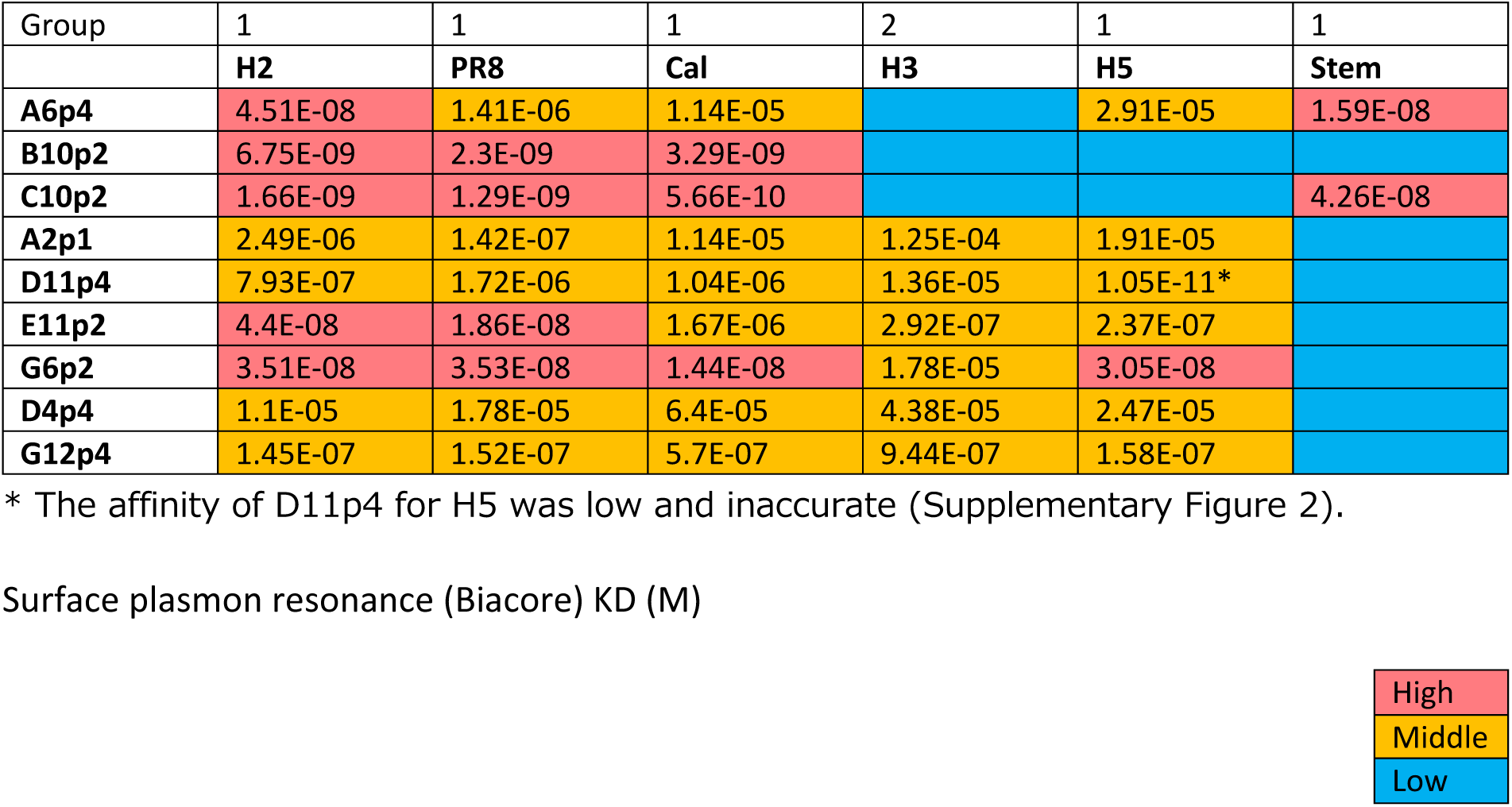
Affinities (K_d_) of the antibodies to six HA antigens from strains A/Okuda/1957(H2N2), termed H2, A/Puerto Rico/8/1934(H1N1), termed PR8, A/California/2009 (X-179A) [H1N1] Pdm09, termed Cal, A/Texas/50/2012 (X-223) [H3N2], termed H3, A/Egypt/N03072/2010(H5N1), termed H5, and A/Brisbane/59/2007 (H1N1), termed Stem, were determined using surface plasmon resonance analysis. H2, PR8, Cal, and H5 belong to group 1, and H3 belongs to group 2.

| Group | 1 | 1 | 1 | 2 | 1 | 1 |
| --- | --- | --- | --- | --- | --- | --- |
|  | H2 | PR8 | Cal | H3 | H5 | Stem |
| A6p4 | 4.51E-08 | 1.41E-06 | 1.14E-05 |  | 2.91E-05 | 1.59E-08 |
| B10p2 | 6.75E-09 | 2.3E-09 | 3.29E-09 |  |  |  |
| C10p2 | 1.66E-09 | 1.29E-09 | 5.66E-10 |  |  | 4.26E-08 |
| A2p1 | 2.49E-06 | 1.42E-07 | 1.14E-05 | 1.25E-04 | 1.91E-05 |  |
| D11p4 | 7.93E-07 | 1.72E-06 | 1.04E-06 | 1.36E-05 | 1.05E-11* |  |
| E11p2 | 4.4E-08 | 1.86E-08 | 1.67E-06 | 2.92E-07 | 2.37E-07 |  |
| G6p2 | 3.51E-08 | 3.53E-08 | 1.44E-08 | 1.78E-05 | 3.05E-08 |  |
| D4p4 | 1.1E-05 | 1.78E-05 | 6.4E-05 | 4.38E-05 | 2.47E-05 |  |
| G12p4 | 1.45E-07 | 1.52E-07 | 5.7E-07 | 9.44E-07 | 1.58E-07 |  |

\* The affinity of D11p4 for H5 was low and inaccurate (Supplementary Figure 2).

Surface plasmon resonance (Biacore) $K_D$ (M)
|  |
| --- |
| High |
| Middle |
| Low |

Furthermore, six antibodies that bound to H3 were categorized into group 2 (23). In addition, we observed that two of the broadly reactive antibodies recognized the stem region of the HA of A/Brisbane/59/2007. A possible scenario for other stem-negative antibodies is that they may either bind to the stem region of other strains or to the head region of HA. We tested whether some of these antibodies shared features with classic broadly neutralizing antibodies. C179, the first broadly neutralizing mouse antibody isolated in 1993 (17), reacts with the stem region of HAs from group 1 influenza virus strains (23). C179 binds to HAs from group 1 influenza viruses, including H1N1, H2N2, and H5N1. A competition assay for the binding of H2 and PR8 revealed that A6p4 competes with C179 (Fig. 3). In addition, our two-dimensional phylogeny map indicated that A6p4 was unique to all clones (Supplementary Fig. 3). Note that we used NSP2 as a negative control antibody, which binds specifically to the HA from A/California/2009 (X-179A) [H1N1] Pdm09 (24). These results demonstrate that we were able to isolate a broadly reactive mAb with an affinity comparable to that of a classic broadly reactive mAb (C179) using our technology.

**Figure 3.**
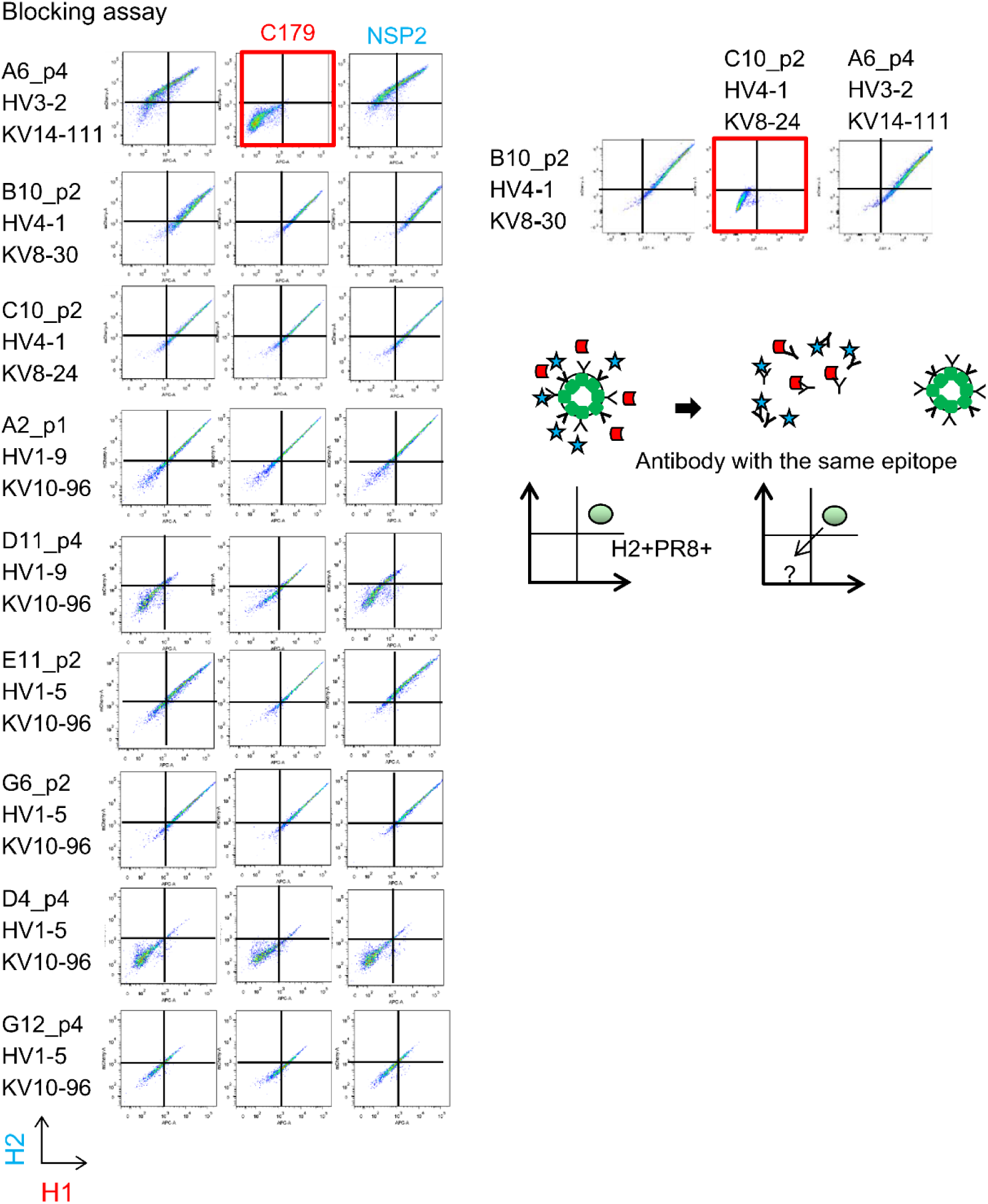
Competition assay of “cross” Ig-transformants using C179, a broadly-reactive mAb binding to the Stem region of HA, and NSP2 (Narita strain-specific mAb), a strain-specific mAb that binds to the head region of HA. On the right panel, two stem-reactive clones (see Table 1) were examined reciprocally with combination of transformant and recombinant antibodies.

## Discussion

The discovery of therapeutic antibodies as well as their demand are increasing (25). Efficiently isolating one biologically significant antibody from the 10^13^ antibody repertoire is challenging. The hybridoma method has long been used for this purpose and remains reliable. However, the fusion rate of B and myeloma cells is < 0.001% in this method; therefore, the number of hybridomas obtained from one individual B cell is limited to several hundred clones, limiting its use in the production of mAbs. As an alternative choice, cloning-based antibody screening has gained popularity, especially for human antibody discovery, because of the lack of a human myeloma cell line similar to that used for murine cell fusion partners. The SPYMEG cell line can generate antigen-specific antibody-producing cells by fusing the murine SP2/0 myeloma cell line and human megakaryoblastic leukemia cell line MEG-01 with human PBMCs (14, 26). In our study, we sorted IgG1 + B cells. Most of those cross-reactive B cells were memory B cells (Supplementary Figure 4). As shown in Supplementary Figure 5, the cloning efficiency exceeded 75%. Our functional screening by selecting strong binders allowed us to narrow down the number of clones to 13 cross-reactive B cells. One of the clones had a strong affinity of K_d_ ≃ 5.66 × 10^­10^ (M). Notably, the generation of antibodies that target regions other than HA cannot be ruled out since the immunization antigen and the detected antigen were the same. However, as shown in Table 1, the cross-reactive antibodies obtained in this study exhibited characteristic binding abilities to each of the six types of HAs. If these were antibodies recognizing His-tag, they would bind to all six types of HA, indicating that these cross-reactive antibodies were no His-specific clones.

In addition to gene expression profiling, NGS facilitates antibody repertoire analysis. However, despite breakthroughs in sequencing technology, the process of linking antibody function and its genes is still long and labor intensive. To overcome this issue, we developed a new antibody screening system to directly link antigen–antibody binding (as a function) with a gene encoding the antibody by expressing its membrane-bound form.

Among conventional cloning-based mAb isolation methods, this system demonstrates several advantages. 1) Membrane Ig expression can link the antigen-binding feature of membrane-expressed Ig, which can be linked to Ig DNA sequence information using our plasmid construct. Similar to the panning procedure used in phage display (27), antigen-binding cells carry the plasmid encoding Ig genes through surface Ig so that the selection process in bulk format enriches the relevant plasmids for use in further experiments. 2) The dual Ig expression vector links heavy- and light-chain genes, which reduces the plasmid preparation time and stock by half. 3) Highly reliable Golden Gate Cloning technology using type IIs restriction enzymes can readily generate plasmid clones. This reduced the time required to generate an Ig plasmid library. 4) The population profile, defined by the fluorescence intensity during flow cytometry, directly reflected the affinity of a clone (Supplementary Figure 6). Lima et al. developed a dual Ig expression vector, similar to the one we report here, to produce secretary antibodies and succeeded in obtaining functional monoclonal antibodies from SARS-CoV-2 infected individuals (28). Collectively, our technology streamlines the isolation of mAbs for therapy and diagnosis.

## Conclusion

Our findings indicate that the developed antibody presentation system facilitates antibody functional analysis and is well suited for the discovery of antibodies important for infectious diseases when combined with conventional NGS-based antibody repertoire analysis. Compared with droplet-based experimental systems, well-based systems are limited in the number of cells they can process. Furthermore, experiments involving infectious bacteria and viruses have imposed limitations on human experimentation. To solve these problems, the automation of experiments will become important in the future.

By combining our screening system with robotic automation of experiments, it will be possible to obtain useful mAbs for various diseases quickly and in large quantities, which has broad implications for the development of vaccines against various diseases.

## Materials and Methods

### Protein purification of influenza HAs (H1 and H2) as antigens

A mixture of 25.7 µg of HA, comprising H1 strain A/Puerto Rico/8/1934 (PR8), H1 strain A/California/7/2009 (X-179A), H2 strain A/Okuda/1957 (H2), H3 strain A/Texas/50/2012 (X-223), or H5 strain A/Egypt/N03072/2010], 1.3 µg of NA (Strain-matched), and 3 µg of BirA-expressing plasmids were transfected into Expi293 cells using the Expifectamine 293 transfection kit (Thermo Fisher Scientific, Waltham, MA, USA) according to the manufacturer’s instructions. The cells were then cultured in Expi293 Expression Medium (Thermo Fisher Scientific) supplemented with D-biotin at a final concentration of 100 µM for *in-vivo* BirA biotinylation in a humidified incubator containing 8% CO_2_ at 37 °C and 125 rpm. If biotinylation was not necessary, transfection was performed without BirA, and cell culture was carried out without D-biotin. Culture supernatants were harvested 3 days post-transfection and filtered through a 0.45-μm filter (Merck Millipore, Burlington, MA, USA). All proteins were purified with a Talon metal affinity resin (Takara Bio USA, Mountain View, CA, USA) and dialyzed against pyrogen-free PBS. The oligomeric state and purity were determined at 25 °C by size-exclusion chromatography on a Superdex 200 10/300GL column (GE Healthcare Technologies, Chicago, IL).

### HA (H1 and H2) protein immunization

Two BALB/c mice were intraperitoneally immunized sequentially, 2 weeks apart, with 15 µg of H1 (PR8) HA, followed by 15 µg of H2 HA protein as an antigen and supplemented with AddaVax adjuvant (InvivoGen, San Diego, CA, USA). Two weeks after the second immunization, the mice were sacrificed, and their splenocytes were used for the experiment. BALB/c mice were purchased from CLEA Japan (CLEA Japan, Inc., Tokyo, Japan). All animal experiments were performed using protocols approved by the Institutional Animal Care and Use Committee (IACUC) of the RIKEN Yokohama Branch.

### Fluorescence-activated cell sorting

Using ACK (Ammonium-Chloride-Potassium) lysing buffer-treated splenocytes, CD43-negative B cells were collected using AutoMACS (Miltenyi Biotec, Inc., Bergisch Gladbach, North Rhine-Westphalia, Germany) and stained with IgG1 (Bv510), non-biotinylated His-tagged purified recombinant His-tagged H1 (PR8) protein, and CD38 (PE-Cy7) on ice for 30 min. After washing three times, the cell suspension was incubated with Alexa Fluor 488-labeled anti-His antibodies on ice for 30 min. After washing three times, biotinylated purified recombinant H2 protein was incubated on ice for 30 min. Furthermore after washing three times, the cell suspension was incubated with Brilliant Violet 421-conjugated streptavidin on ice for 3 min. We used 7-AAD for excluding dead cells. Next, the B cells were subjected to single-cell sorting using a BD FACSAria III (BD Biosciences, Franklin Lakes, NJ, USA) and collected into 96-well plates pre-loaded with lysis buffer. The single cells in the lysis buffer were immediately snap-frozen on powdered dry ice and stored at ­80 °C.

### Amplification of a paired B-cell repertoire amplicon from a single cell

Single cells were dropped into pre-loaded plates containing 4 µL/well of lysis buffer (2 U/µL RNase inhibitor, 0.5× PBS, 10 mM DTT) and 1 µL of 10 µM Oligo(dT)18 Primer. Cell lysates were incubated at 70 °C for 90 s, followed by incubation at 35 °C for 15 s, and then placed on ice for 2 min. For first-strand cDNA synthesis, 5 µL of RT mix (2× PCR buffer, 5 mM DTT, 2 mM dNTP Mix, 10 U RNase inhibitor, 100 U SuperScript III Reverse Transcriptase) was added to the cell lysate and incubated consecutively at 35 °C for 5 min, 45 °C for 20 min, 70 °C for 10 min, and 4 °C for chilling. For primer digestion, 5 µL of the primer digestion mix (1× ExoI Buffer, 7.5 U Exonuclease I) was prepared and incubated at 37 °C for 30 min, 80 °C for 20 min, and 4 °C for chilling. For the poly-A tailing reaction, 5 µL of the poly-A tailing reaction mix (4× PCR buffer, 5 mM dATP, 0.1 U RNase H, 60 U Terminal transferase) was prepared and incubated at 37 °C for 50 s, 65 °C for 10 min, and 4 °C for chilling. For the 2nd strand cDNA synthesis, 25 µL of the amplification mix (1× KAPA HiFi HotStart ReadyMix, 400 nM SMART_dT primer, 10 µL of poly-A tailing product, 1.5 µL of nuclease-free H_2_O) was prepared and incubated at 95 °C for 3 min, 40 °C for 1 min, 65 °C for 10 min, and 72 °C for 5 min and chilled. For the cDNA amplification using an Ig-specific outer primer set, 20 µL of the outer amplification mix (1× KOD FX Neo buffer, 400 µM dNTPs, 1× UPM, 300 nM outer primer, 3.5 µL of the SMART product, 0.5 U KOD FX Neo polymerase) was prepared and incubated at 94 °C for 2 min, followed by 25 cycles at 98 °C for 10 s, 60 °C for 30 s, and 68 °C for 45 s, and then chilled. For cDNA amplification with an Ig-specific inner primer set, 20 µL of the inner amplification mix (1× KOD FX Neo buffer, 400 µM dNTPs, 500 nM 5’ matrix primer mix, 500 nM 3’ matrix primer mix, 3.5 µL of the outer cDNA amplified product, 0.5 U KOD FX Neo polymerase) was prepared and incubated at 94 °C for 2 min, followed by 25 cycles at 98 °C for 10 s, 60 °C for 30 s, and 68 °C for 45 s and then chilled. Finally, 3 µL of the amplified inner cDNA product was used for further plate indexing by amplification. Then, 20 µL of the plate-indexing amplification mix (1× KOD FX Neo buffer, 400 µM dNTPs; 500 nM i5-Fwd or i7-Fwd, 500 nM i7-Rev or i5-Rev; 2 µL of the inner cDNA amplified product; 0.5 U KOD FX Neo polymerase) was prepared and incubated at 94 °C for 2 min, followed by 25 cycles at 98 °C for 10 s, 60 °C for 30 s, and 68 °C for 45 s, and then chilled. All primers used are listed in Supplementary Tables 1 and 2.

### Paired-end Illumina sequencing strategy and analysis

The libraries were sequenced on a MiSeq instrument (Illumina, San Diego, CA, USA) following the manufacturer’s protocol. As the sequencing length was limited to 2 × 300 bp, we used the following strategy. Amplicons in both orientations were generated using an Illumina adapter sequence. Then, asymmetric 400+100 nt paired-end sequencing was performed to obtain high-quality 400 + 400 nt paired-end sequences of the original cDNA molecule. This approach provided a sufficient sequencing length to capture extra-long Ig variants. We followed the method by Turchaninova et al. (29). Only reads that met the following four conditions were used as the resulting Ig sequences: 1) reads with a length of 260 nt or more; 2) cumulative clustered reads representing the top 5% of the entire reads; 3) 5’- and 3’-reads overlapping a length of 20 bases or more; and 4) a total length of 460 ± 50 bp for heavy chain or 390 ± 40 bp for light chain. All resulting sequence data were analyzed using IMGT/High V-QUEST (http://www.imgt.org/IMGT_vquest/vquest) for annotation (30). Using the sequences and annotations, Ig database construction and visualization of the Ig repertoire were performed using the in-house BONSCI software.

### Construction of a mouse IgG/IgK expression plasmid

The BsaI restriction sites were inserted into the paired B-cell repertoire amplicon from a single cell using PCR. The destination vector contained the IL6 signal peptide, EF1a promoter, ccdB gene, and mouse IgK Fc region. The donor vector contained the mouse IgG1 Fc region, the *Venus* gene, and the EF1a promoter. The paired B-cell repertoire amplicons, destination vector, and donor vector containing BsaI restriction sites were assembled using our assembly method. For this step, 10 μL of the assembly mix (1× T4 DNA ligase buffer, 1× BSA, 1 U BsaI restriction enzyme, 40 U T4 DNA ligase, 100 ng heavy chain amplicon, 100 ng light chain amplicon, 100 ng destination vector, and 100 ng donor vector) was prepared and incubated for 25 cycles at 37 °C for 3 min, 16 °C for 4 min, 50 °C for 5 min, and 80 °C for 5 min, and then chilled. The antibody sequence that entered the construct was fused to the *Venus* sequence and expressed in membrane form.

### Antibody display on FreeStyle 293 cultured cells

One microgram of antibody-expressing plasmid was transfected into 1 x 10^6^ FreeStyle 293 cells in 1-mL culture using the 293fectin Transfection Reagent (Thermo Fisher Scientific) according to the manufacturer’s instructions and cultured in FreeStyle 293 Expression Medium (Thermo Fisher Scientific) in a humidified incubator with 8% CO_2_ at 37 °C and 125 rpm. Antibodies were displayed on the surface of cultured cells, which enabled the determination of antigen specificity. Furthermore, the antibody-display cells were tested for binding activity with Alexa647-labeled H1 and Alexa568-labeled H2 (Alexa Fluor™ Antibody Labeling Kits, ThermoFisher Scientific) using a BD FACSAria III (BD Biosciences).

### Antibody production

To obtain a large amount of secretory antibodies, PCR fragments from the antibody-expressing plasmid were cloned into the pcDNA3.4-mIgG1 or pcDNA3.4-kappa vectors. A mixture of 15 μg of pcDNA3.4-V-gene-mIgG1 vector and 15 μg of pcDNA3.4-V-gene-kappa vector was transfected into Expi293 cells using the ExpiFectamine 293 Transfection Kit (Thermo Fisher Scientific), according to the manufacturer’s instructions, and cultured in Expi293 Expression Medium (Thermo Fisher Scientific) in a humidified 8% CO_2_ incubator at 37 °C and 125 rpm. Then, 5 days post-transfection, the culture supernatants were harvested, and all proteins were purified with PureSpeed IMAC resin (Mettler Toledo, Columbus, OH, USA), according to the manufacturer’s instructions.

### Surface plasmon resonance

Kinetic analyses were performed at 25 °C using a BIAcore 3000 machine (GE Healthcare Technologies). A2p1, B10p2, C10p2, E11p2, G6p2, A6p4, D4p4, D11p4, or G12p4 antibodies were immobilized on a CM5 sensor chip (GE Healthcare Technologies) with an amine-coupling kit according to the manufacturer’s instructions. HA probes were serially diluted at five different concentrations and injected at a flow rate of 30 µL/min for 3 min with a dissociation time of 7 min in HBS–EP buffer (10 mM HEPES, pH 7.4, 150 mM NaCl, 3.4 mM EDTA, and 0.005% Surfactant P20). The chip was regenerated using 10 mM glycine (pH 2.5) as an HA probe.

## Data access

All raw and processed sequencing data generated in this study have been submitted to the NCBI Gene Expression Omnibus (GEO; https://www.ncbi.nlm.nih.gov/geo/) under accession number GSE140720.

## Acknowledgements

We thank Mr. Atsuo Kobayashi, Ms. Chieko Okamura, Dr. Seok-Won Kim, Mr. Maxime Hebrard, Dr. Todd D Taylor, and the RIKEN IMS Genome Platform for their technical support, Dr. Atsushi Miyawaki for gifting the *Venus* gene, and Dr. Christopher Nicholas for critical reading and English editing of the manuscript. We would like to thank Editage (www.editage.jp) for English language editing.

## Author contributions

Conceptualization: Takashi Watanabe, Hidehiro Fukuyama; Data curation: Yoshiki Mochizuki, Naveen Kumar; Formal analysis: Yoshiki Mochizuki, Naveen Kumar; Funding acquisition: Osamu Ohara, Tomohiro Kurosaki, Takashi Watanabe, Hidehiro Fukuyama; Investigation: Hikaru Hata, Fumie Yokoyama, Tomoko Hasegawa; Methodology: Takashi Watanabe, Hidehiro Fukuyama; Project administration: Takashi Watanabe, Hidehiro Fukuyama; Resources: Takashi Watanabe, Hidehiro Fukuyama; Software: Yoshiki Mochizuki, Naveen Kumar; Supervision: Osamu Ohara, Tomohiro Kurosaki; Validation: Takashi Watanabe, Hidehiro Fukuyama; Visualization: Takashi Watanabe, Hidehiro Fukuyama; Writing – original draft: Takashi Watanabe, Hidehiro Fukuyama; Writing– review & editing: Takashi Watanabe, Hidehiro Fukuyama.

## Supplementary Tables

**Supplementary Table 1.**
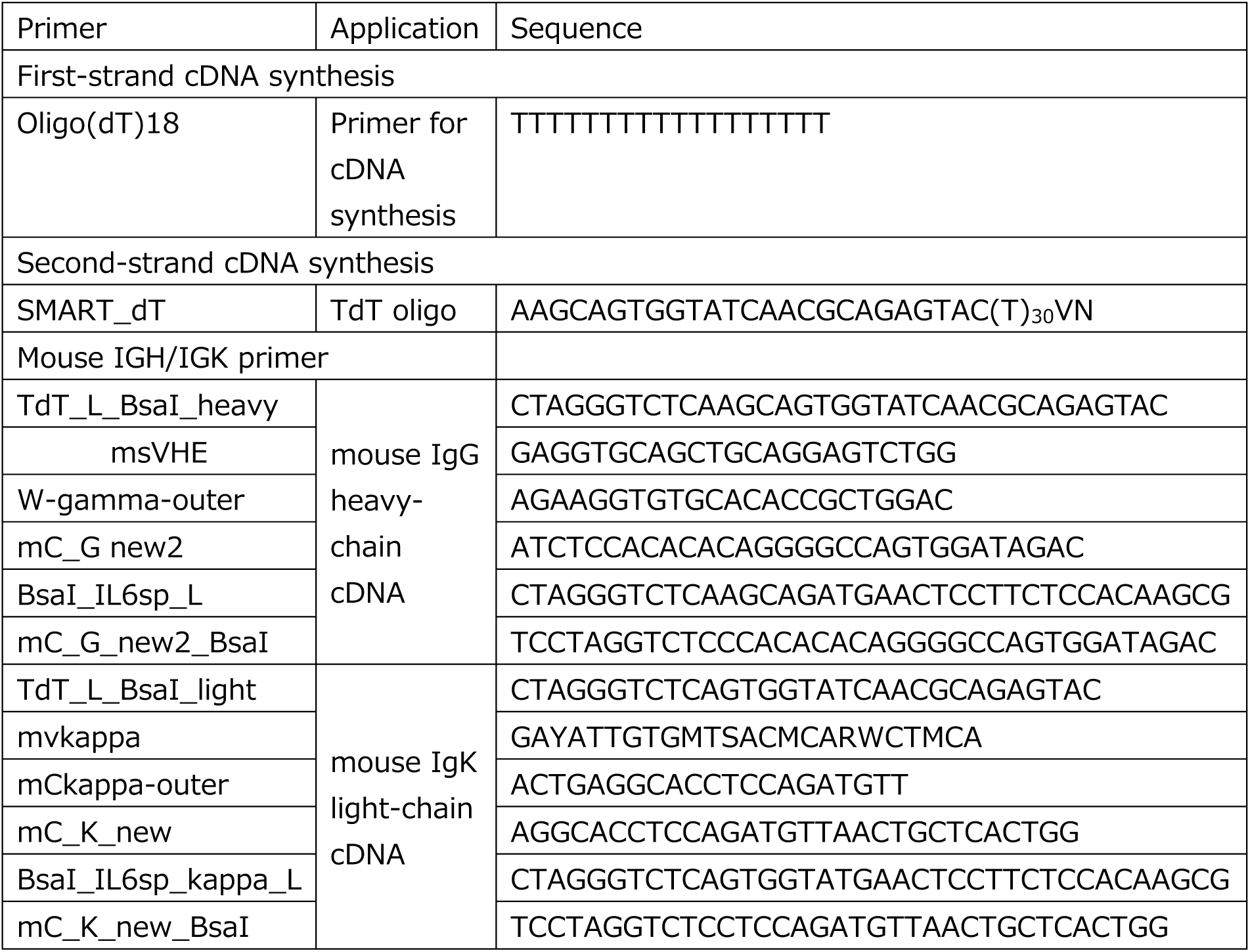
Oligonucleotides.

**Supplementary Table 2.**
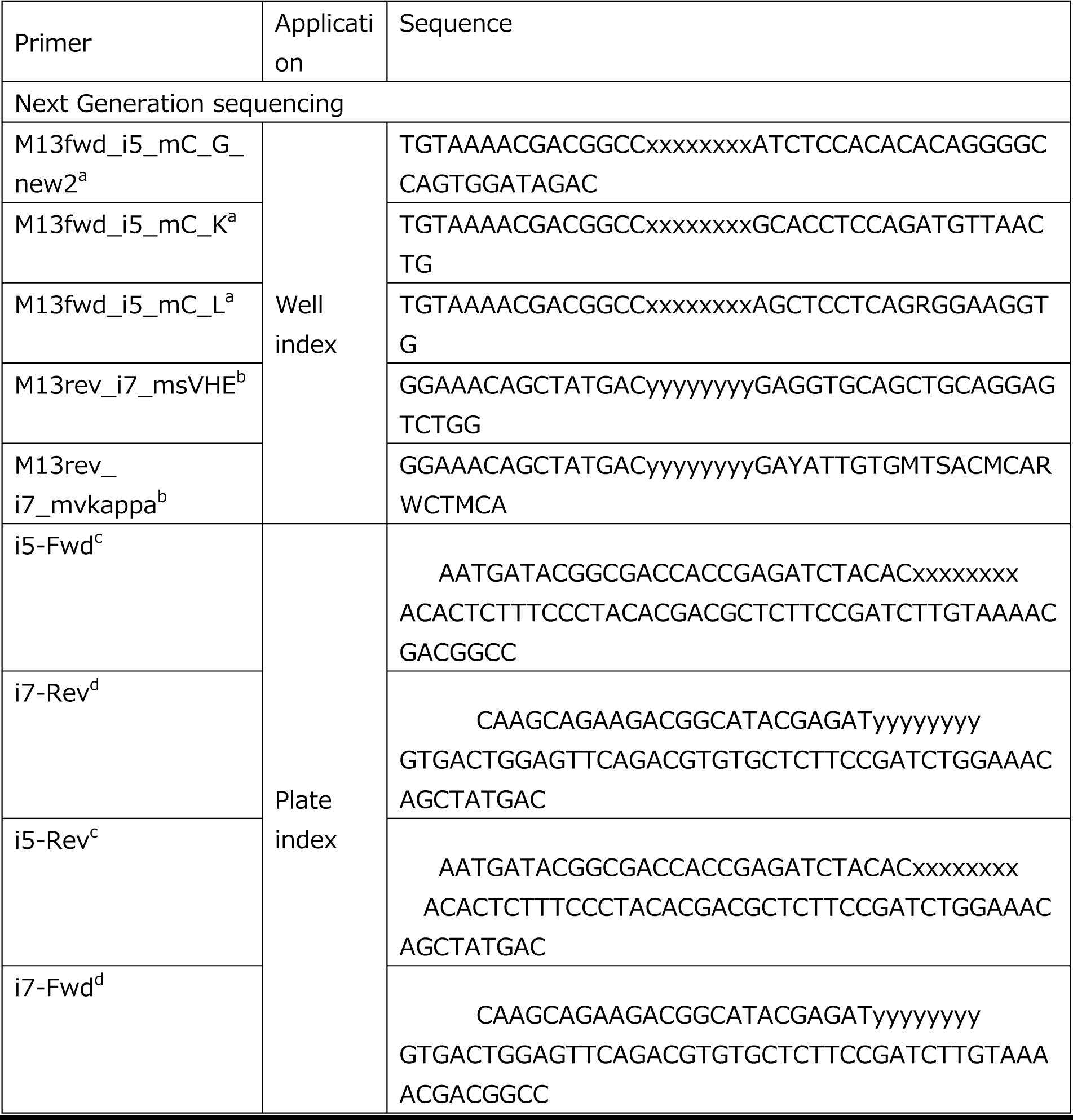
Oligonucleotides.

## Supplementary Figures

**Supplementary Figure 1.**
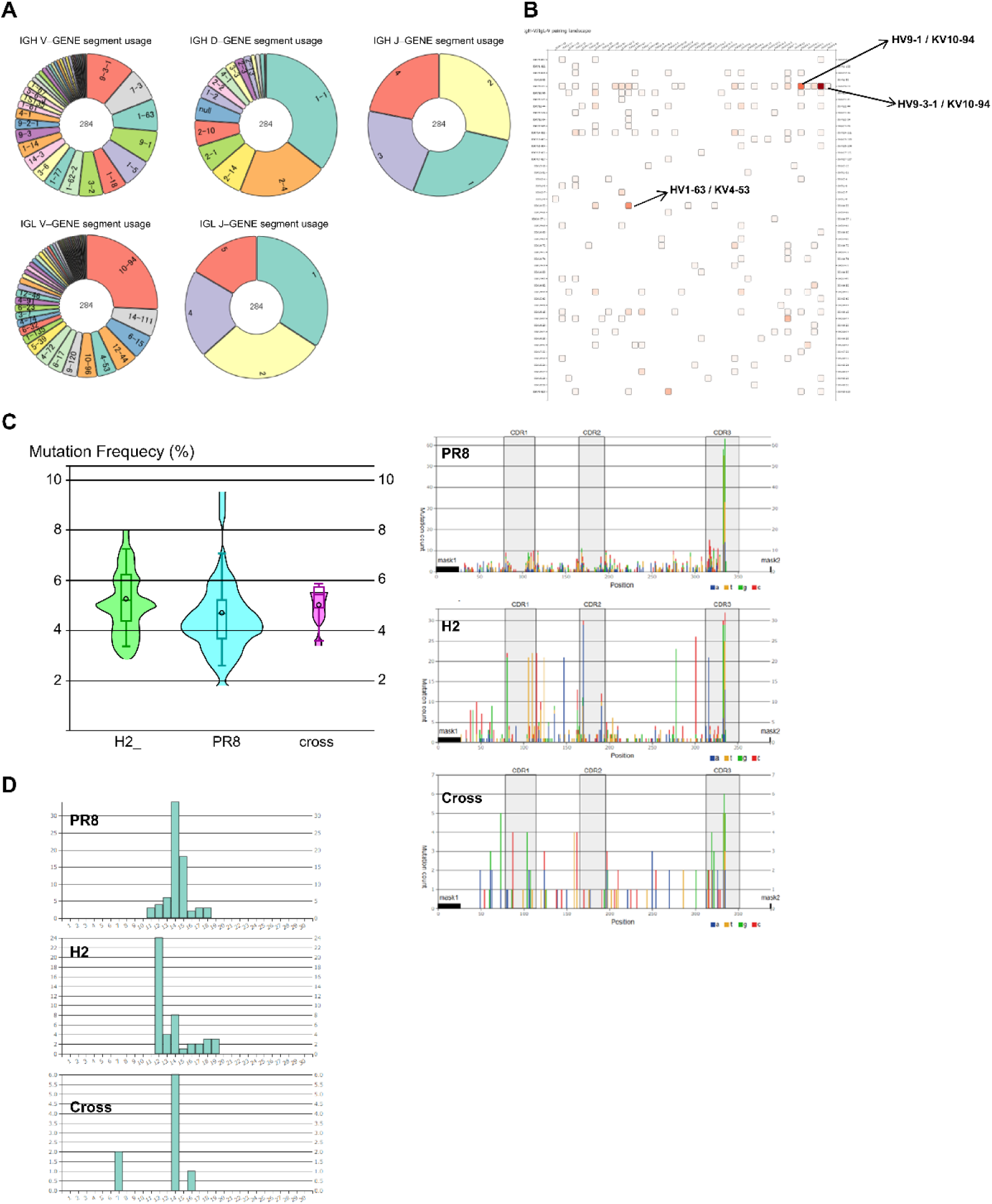
The Ig repertoires are visualized using the in-house software. Overview of heavy (V-D-J) and light (V-J) chain usages (**A**), and the repertoire clonality (**B**) is shown. (**C**) The mutation rates and their positions of heavy chains in three cell populations (PR8+, H2+, PR8+H2+) are compared. (**D**) The lengths of CDR3 of heavy chains of three cell populations (PR8+, H2+, PR8+H2+) are compared.

**Supplementary Figure 2.**
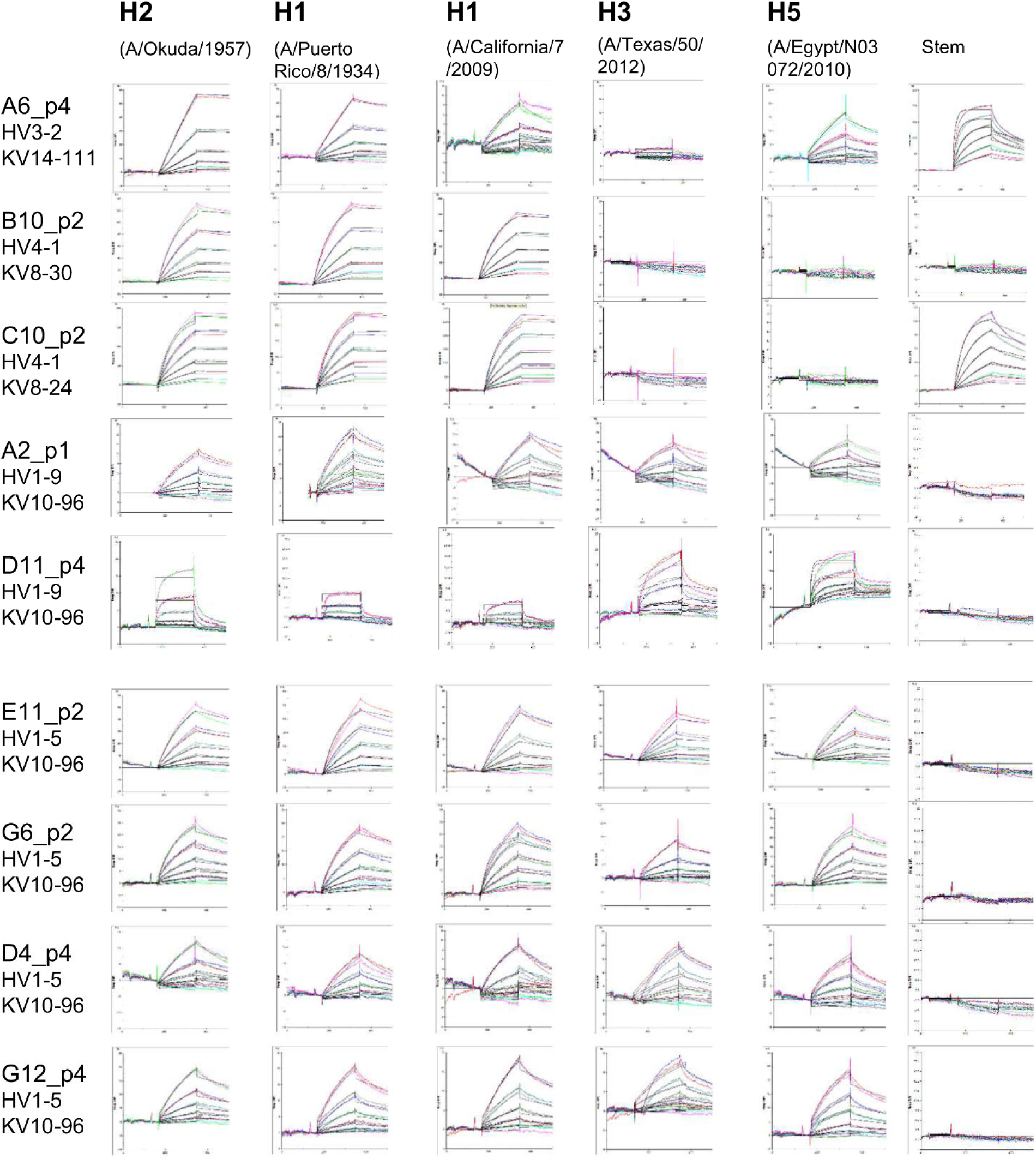
SPR profiles of kinetics analysis of nine clones, A6p4, B10p2, C10p2, A2p1, D11p4, E11p2, G6p2, D4p4, and G12p4 using a BIAcore 3000 machine (GE Healthcare).

**Supplementary Figure 3.**
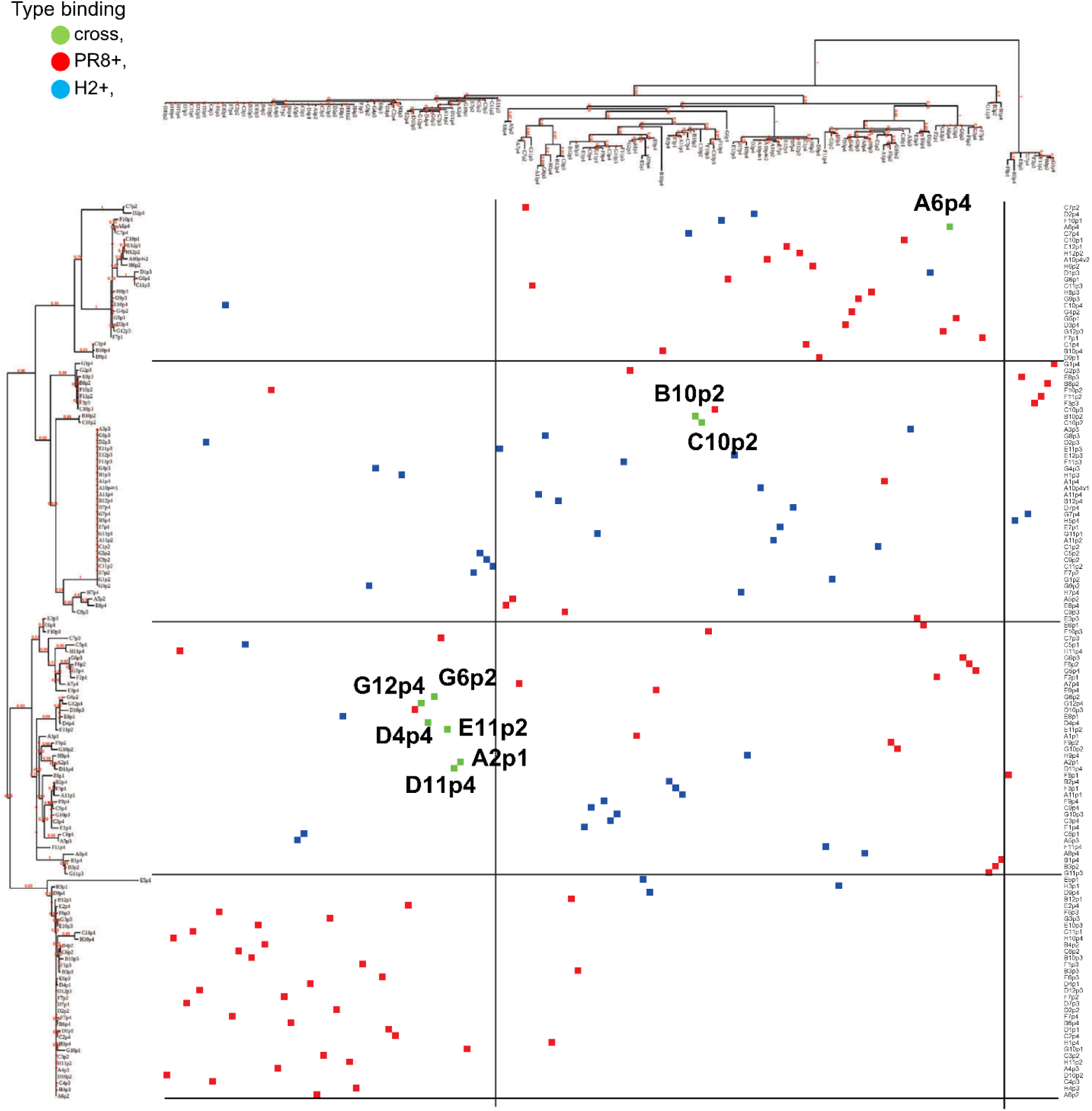
Two hundred and eighty-four clones are plotted in a two-dimensional map of phylogeny trees for heavy and light chain genes. Nine mAb clones are overlaid in green. Six “cross” clones are located in a major cluster. B10p2 and C10p2 are co-localized and distant from the cluster. A6p4 is uniquely located in the map.

**Supplementary Figure 4.**
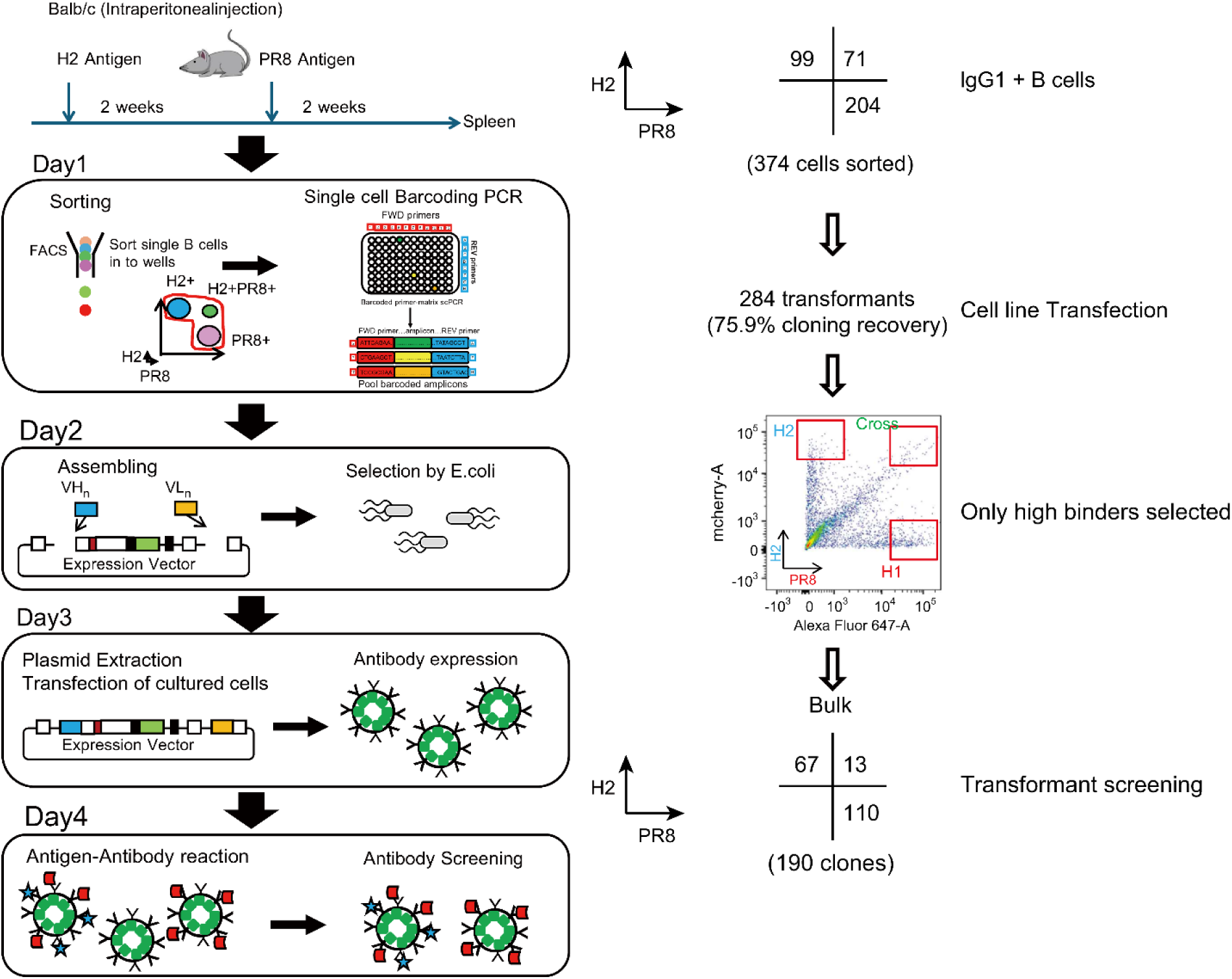
Progression Flowchart. The process for efficiently isolating influenza cross-reactive antibodies from mouse germinal center B cells with high affinity is shown. Experiment outcome flow numbers of clones in each step are indicated.

**Supplementary Figure 5.**
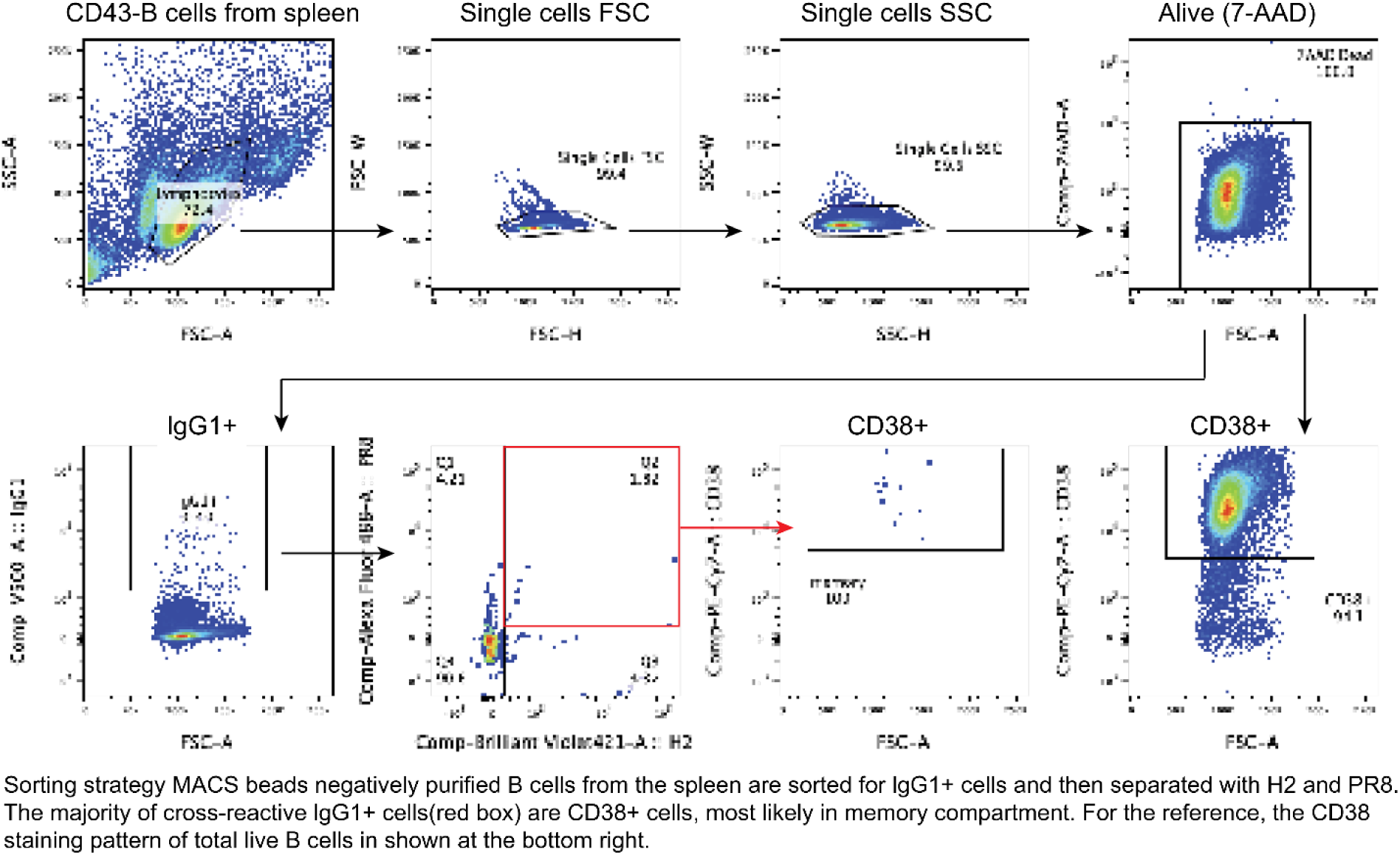
Sorting strategy. CD43 MACS B cells from the spleen purified using negatively charged beads were sorted for IgG1+ cells and then separated into H2 and PR8. The majority of cross-reactive IgG1+ cells (red box) were CD38+ cells, most likely in the memory compartment. For reference, the CD38 staining pattern of total live B cells is shown at the bottom right.

**Supplementary Figure 6.**
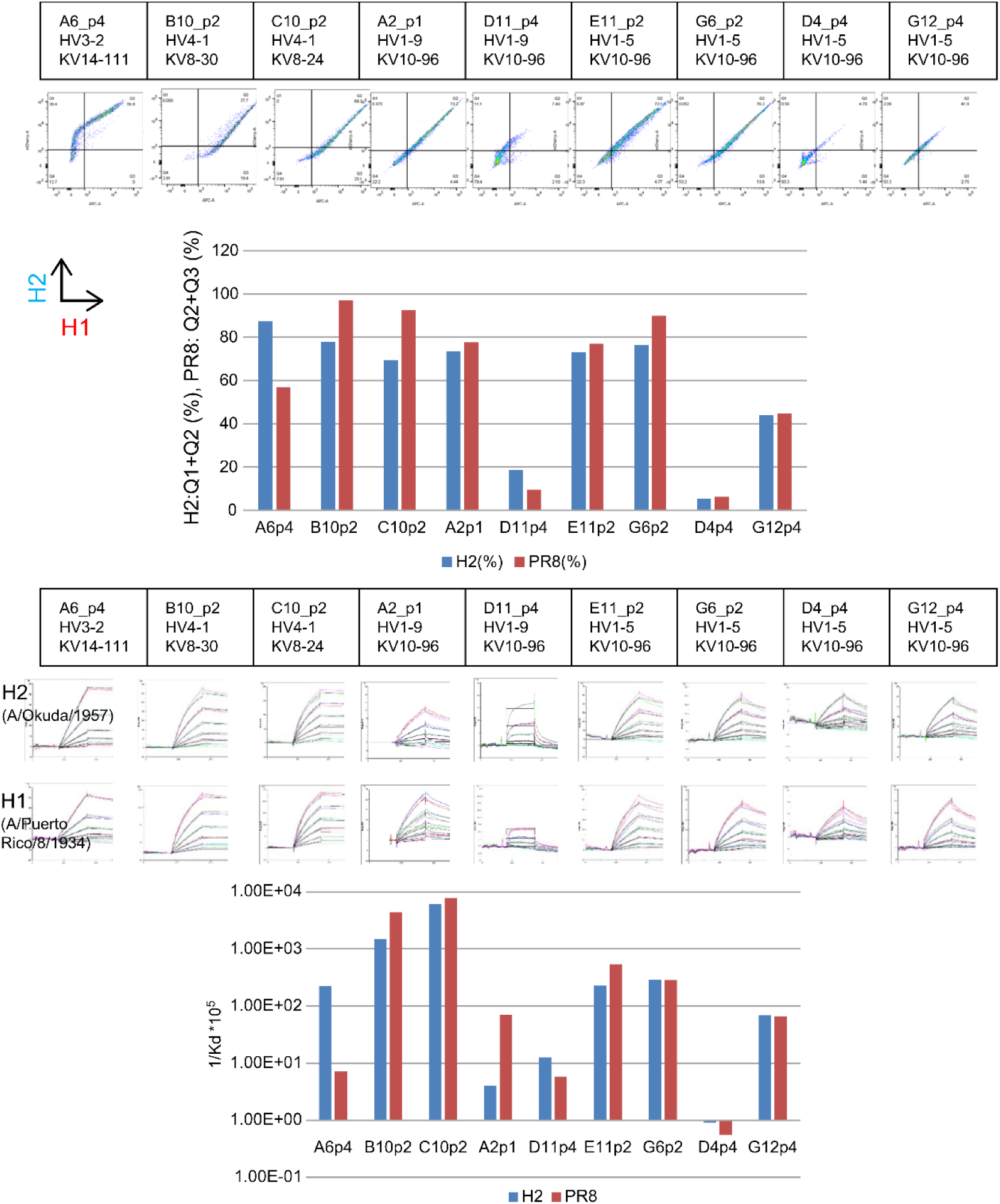
The affinity of antigen-antibody binding (in this case, probe and membrane-bound antibody expressing cell) can be inferred from the population shift (flow cytometry analysis).

## References

1. Balsitis, S. J., K. L. Williams, et al. (2010). Lethal antibody enhancement of dengue disease in mice is prevented by Fc modification. PLoS Pathog., 6(2): e1000790.

2. Fang, X. T., D. Sehlin, et al. (2017). Efficient and inexpensive transient expression of multispecific multivalent antibodies in Expi293 cells. Biol. Proced. Online, 19(1): 1–9.

3. Köhler, G. and C. Milstein (1975). Continuous cultures of fused cells secreting antibody of predefined specificity. Nature, 256(5517): 495–497.

4. Traggiai, E., S. Becker, et al. (2004). An efficient method to make human monoclonal antibodies from memory B cells: potent neutralization of SARS coronavirus. Nat. Med., 10(8): 871–875.

5. Kwakkenbos, M. J., S. A. Diehl, et al. (2010). Generation of stable monoclonal antibody-Producing B cell receptor-positive human memory B cells by genetic programming. Nat. Med., 16(1): 123–128.

6. Wrammert, J., K. Smith, et al. (2008). Rapid cloning of high-affinity human monoclonal antibodies against influenza virus. Nature, 453(7195): 667–671.

7. Meijer, P. J., P. S. Andersen, et al. (2006). Isolation of human antibody repertoires with preservation of the natural heavy and light chain pairing. J. Mol. Biol., 358(3): 764–772.

8. Kuraoka, M., A. G. Schmidt, et al. (2016). Complex antigens drive permissive clonal selection in germinal centers. Immunity, 44(3): 542–552.

9. Clackson, T., H. R. Hoogenboom, et al. (1991). Making antibody fragments using phage display libraries. Nature, 352(6336): 624–628.

10. Feldhaus, M. J., R. W. Siegel, et al. (2003). Flow-cytometric isolation of human antibodies from a nonimmune Saccharomyces cerevisiae surface display library. Nat. Biotechnol., 21(2): 163–170.

11. Harvey, B. R., G. Georgiou, et al. (2004). Anchored periplasmic expression, a versatile technology for the isolation of high-affinity antibodies from Escherichia coli-expressed libraries. Proc. Natl. Acad. Sci. U. S. A., 101(25): 9193–9198.

12. Schaffitzel, C., J. Hanes, et al. (1999). Ribosome display: an in vitro method for selection and evolution of antibodies from libraries. J. Immunol. Methods, 231(1–2): 119–135.

13. Mazor, Y., T. Van Blarcom, et al. (2007). Isolation of engineered, full-length antibodies from libraries expressed in Escherichia coli. Nat. Biotechnol., 25(5): 563–565.

14. Kubota-Koketsu, R., H. Mizuta, et al. (2009). Broad neutralizing human monoclonal antibodies against influenza virus from vaccinated healthy donors. Biochem. Biophys. Res. Commun., 387(1): 180–185.

15. Setliff, I., A. R. Shiakolas, et al. (2019). High-throughput mapping of B cell receptor sequences to antigen specificity. Cell, 179(7): 1636–1646. e15.

16. Kirchmaier, S., K. Lust, et al. (2013). Golden GATEway cloning--a combinatorial approach to generate fusion and recombination constructs. PloS One, 8(10): e76117.

17. Okuno, Y., Y. Isegawa, et al. (1993). A common neutralizing epitope conserved between the hemagglutinins of influenza A virus H1 and H2 strains. J. Virol., 67(5): 2552–2558.

18. Ekiert, D. C., A. K. Kashyap, et al. (2012). Cross-neutralization of influenza A viruses mediated by a single antibody loop. Nature, 489(7417): 526–532.

19. Impagliazzo, A., F. Milder, et al. (2015). A stable trimeric influenza hemagglutinin stem as a broadly protective immunogen. Science, 349(6254): 1301–1306.

20. Nagai, T., K. Ibata, et al. (2002). A variant of yellow fluorescent protein with fast and efficient maturation for cell-biological applications. Nat. Biotechnol., 20(1): 87–90.

21. D’Angelo S, Ferrara F, et al. (2018). Many Routes to an Antibody Heavy-Chain CDR3: Necessary, Yet Insufficient, for Specific Binding. Front. Immunol., 9(395)

22. Xu, J. L. and M. M. Davis (2000). Diversity in the CDR3 region of VH is sufficient for most antibody specificities. Immunity, 13(1): 37–45.

23. Gamblin, S. J., and J.J. Skehel. (2010). Influenza hemagglutinin and neuraminidase membrane glycoproteins. J. Biol. Chem., 285(37): 28403–28409.

24. Adachi, Y., T. Onodera, et al. (2015). Distinct germinal center selection at local sites shapes memory B cell response to viral escape. J. Exp. Med., 212 (10): 1709–1723.

25. Kaplon H, Crescioli S, et al. (2023). Antibodies to watch in 2023. Mabs, 15(1): 2153410

26. Soni P, A Yasuhara, et al. (2018). Evaluation of the fusion partner cell line SPYMEG for obtaining human monoclonal antibodies against influenza B virus. J. Vet. Med. Sci., 80(6): 1020–1024

27. Petropoulos, K., (2012) Phage display. Methods Mol. Biol., 901: 33–51.

28. Lima, N. S., M. Musayev, et al. (2022). Primary exposure to SARS-CoV-2 variants elicits convergent epitope specificities, immunoglobulin V gene usage and public B cell clones. Nat. Commun., 13(1): 7733.

29. Turchaninova, M. A., A. Davydov, et al. (2016). High-quality full-length immunoglobulin profiling with unique molecular barcoding. Nat. Protoc., 11(9): 1599– 1616.

30. Alamyar, E., P. Duroux, et al. (2012). IMGT® tools for the nucleotide analysis of immunoglobulin (IG) and T cell receptor (TR) V-(D)-J repertoires, polymorphisms, and IG mutations: IMGT/V-QUEST and IMGT/HighV-QUEST for NGS. Methods Mol. Biol., 882: 569–604.

